# Several multiple sequence alignment perturbation methods enhance AlphaFold3 sampling of alternative protein states

**DOI:** 10.64898/2026.04.02.716037

**Authors:** Samuel Eriksson Lidbrink, Ivan Nissen, Jonathan Kenichi Ahrlind, Rebecca J Howard, Erik Lindahl

## Abstract

Protein function often involves multiple conformational states. Several multiple sequence alignment-perturbing strategies, including stochastic subsampling, clustering, and column masking, have been shown to enhance AlphaFold2 (AF2) sampling of alternative protein states. Here, we evaluate these strategies on AlphaFold3 (AF3) and compare their performance with the BioEmu Boltzmann sampling model on 107 proteins with multiple experimentally solved conformational states. We find that unperturbed AF3 samples alternative states with significantly higher TM-scores compared to AF2 and comparable to BioEmu. In particular, all MSA perturbation methods improve AF3 sampling at a statistically significant level, improving the top 1% TM-score by at least 0.05 in approximately 20% of cases each, while rarely worsening the performance. Furthermore, we find that different choices of amino acid masks can improve column-masked AF3 sampling for specific targets. Our results highlight how MSA perturbations remain relevant in AF3, providing a useful tool for understanding dynamic biological processes.

## Introduction

Protein dynamics are crucial to many biological functions such as enzyme catalysis, receptor signaling and molecular transport. Determining the 3D-structures of the proteins in all their functionally relevant states would provide profound insight into how the protein functions, and would for example provide a basis for rational drug design. For several proteins there are atomic-resolution structures captured in different conformations [1–5], but in many cases critical states remain elusive. For these, in-silico predictions of the protein conformational states can be useful, e.g. to simplify model building by providing good starting structures. It is also an important source of candidate structures where high-resolution experimental data is lacking, which is especially useful when combined with functional and/or low-resolution experimental data [6, 7].

AI-based methods such as AlphaFold2 (AF2) [8] can provide predictions of the native state with very high accuracy, but usually only predict a single conformational state per protein [9–11]. It has however been shown that perturbations of the input multiple sequence alignment (MSA) allow AF2 to sample additional conformational states for several systems. This has been done e.g. by stochastically subsampling the MSA depth [12], by clustering rows in sequence space [13], or by stochastically masking different columns in the alignment [14] (Figure 1A). The more recent model AlphaFold3 (AF3) [15] expands AF2’s predictive abilities. Since AF3 employs a diffusion model, it has a theoretical ability to natively sample a probability distribution of different states. A recent study has, however, shown that MSA column masking still may improve the sampling of multiple conformational states [16].

**Fig. 1.**
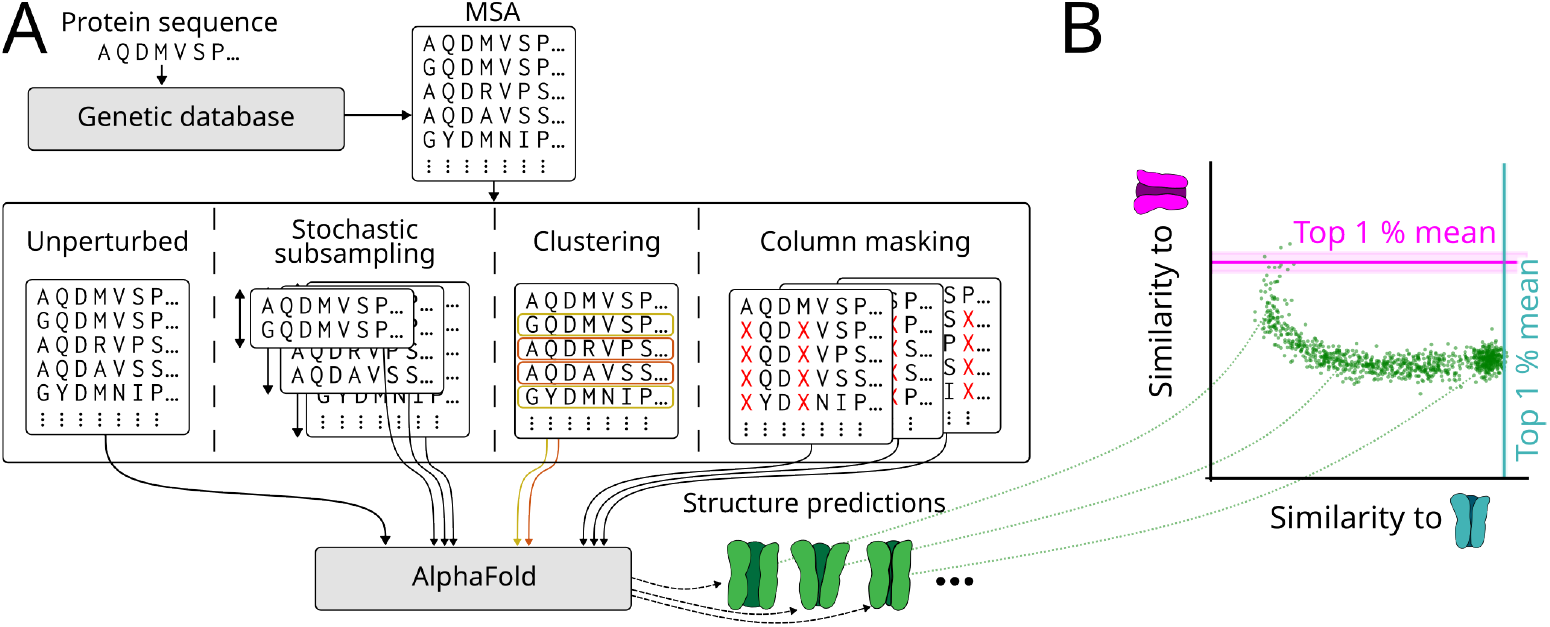
The MSA can be perturbed in different ways with the goal of enhancing AlphaFold sampling of multiple protein states. (A) Overview of the different perturbing methods. The alignment is perturbed either by stochastically subsampling its depth, by clustering the sequences, or by stochastically masking columns. (B) Conceptual diversity plot of the similarity between the structure predictions and experimental references. The ability of a given method to sample a specific state can be benchmarked using the average similarity score for the 1% most similar predictions to a given reference.

Here, we seek to compare the extent to which different MSA perturbation techniques improve AF3 sampling of multiple conformational states, and further compare these methods with the BioEmu model designed to natively sample Boltzmann distributions of the protein conformational landscapes[17]. To do this, we benchmark all these methods on a dataset of more than 100 proteins with experimentally solved structures in multiple conformations. We also explore how the choice of amino acid column mask impacts the sampling. Our results provide insight into the usefulness of the MSA perturbations in AF3, and we contextualize our findings by detailing three specific biological examples.

## Results

### MSA perturbations enhance AF3 sampling of alternative protein states

We examined the effect of three different types of MSA perturbations on AF3 sampling of multiple states: stochastic subsampling, clustering, and column masking (Figure 1A). Given an input MSA, AF3 stochastically samples a subset of the sequences for each recycling iteration to use in its prediction pipeline [15]. In the stochastic subsampling method, the number of input sequences is decreased, which forces AF3 to use shallower MSAs that likely reduces the signal of the dominant state and consequently improves signal-to-noise for alternative states. Conversely, when using the clustering method, the MSA is first clustered in sequence space, and each cluster provided to AF3 separately, where different clusters may carry distinct coevolutionary information. For the column masking method, a stochastically chosen subset of the MSA columns (corresponding to residue positions) is masked with the unknown amino acid X, potentially reducing the coevolutionary signal corresponding to an otherwise dominant state in favor of alternative states. Details of the implementations are provided in the Methods section. We compared these methods to unperturbed AF3, as well as AF2 and BioEmu, all using the same MSA as input.

To assess how well the different methods could sample multiple states of proteins, we examined their ability to sample both experimental structures for 107 proteins from a range of different datasets (see Methods). For each protein and method, at least 1000 protein structures were generated and their similarity assessed against corresponding experimental structures by computing the template modeling score (TM-score) [18], using TM-align [19] evaluated on C*α*-atoms. The TM-score provides a normalized measure of model similarity that is robust to local flexibility and is designed to be independent of protein size, with a score of 1 indicating a perfect match between model and reference [18]. Since the methods may predict multiple different conformational states, we compared the methods by examining the mean of the top 1% TM-scores per reference structure (Figure 1B). Below, we refer to the experimental structure with the higher/lower mean top 1% TM-score as the preferred/alternative state, classified for each method and protein.

Overall, we found that both the alternative and preferred states were sampled with significantly higher top 1% TM-scores in unperturbed AF3 compared to AF2 (using the Wilcoxon signed-rank test for statistical significance) (Figure 2A, Supplementary Figure 1A). MSA column masking, clustering and stochastic subsampling all significantly increased these scores for the alternative states (Figure 2A). We saw a similar improvement when we compared the alternative states per fixed latent space in AF3 (i.e. a given random seed per protein and model) (Supplementary Figure 1B). This indicates that they improve AlphaFold3’s ability to generate a single latent space from which both experimental states can be sampled, and do not require different states to be sampled from distinct latent spaces. Column masking and stochastic subsampling also improved the scores of the preferred states at a statistically significant level, but - notably - not clustering (Supplementary Figure 1A). Out of the total 107 proteins, column masking, clustering, and stochastic subsampling improved/worsened the mean top 1% TM-score when compared to unperturbed AF3 by at least ΔTM ≥ 0.05 in 23/1, 18/2 and 17/0 cases, respectively, for the alternative state, and in 2/0, 3/3 and 0/0 cases, respectively, for the preferred state (Supplementary Figure 2).

**Fig. 2.**
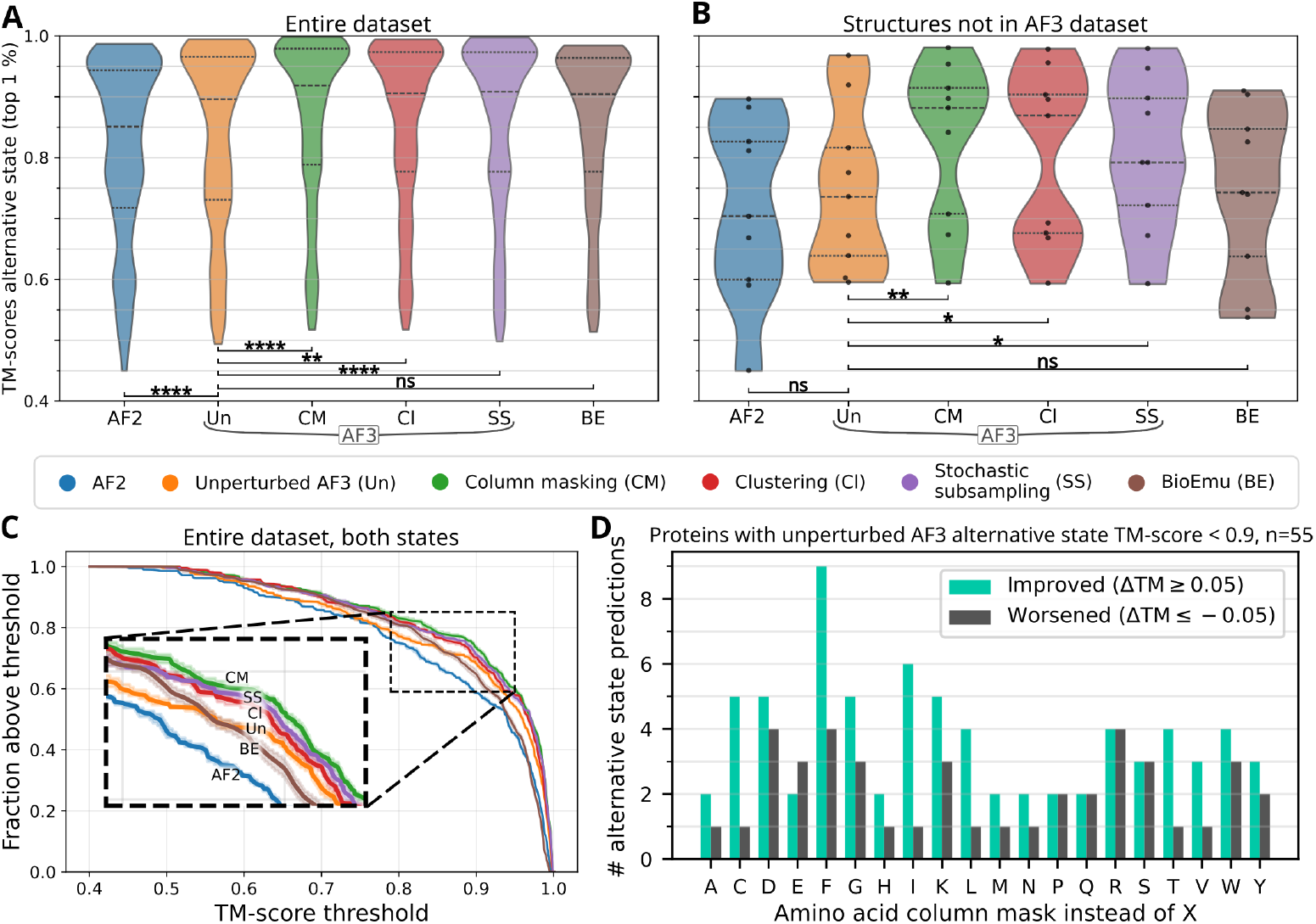
MSA perturbations enhance AF3 sampling of alternative protein states. (A, B) Violin plots showing the distributions of the mean top 1% TM-scores of the alternative state per protein across the different methods for (A) the whole dataset of 107 proteins and (B) the subset of 9 proteins with both experimental structures released after the AlphaFold3 training date cutoff. Statistical significance is assessed using the Wilcoxon Signed-Rank Test (*: *p <* 0.05, **: *p <* 0.01, ***: *p<* 0.001, ****: *p<* 0.0001). Dashed lines indicate quartiles. (C) Fraction of all states with mean top 1% TM scores above different TM-score thresholds, for the various methods across both states for all of the 107 proteins. Shaded regions indicate 95% confidence intervals estimated by bootstrapping the sampled conformations, reflecting the inherent sampling uncertainty. (D) Number of alternative state predictions where the mean top 1% TM-score either improved (teal) or deteriorated (grey) by at least ΔTM ≥ 0.05 when using a specific amino acid column mask instead of X in column-masked AlphaFold3. The amino acid mask variations were sampled for a total of 55 proteins where unperturbed AlphaFold3 sampled at least one state with a mean top 1% TM-score below 0.9.

BioEmu is interesting since the method is trained specifically to perform statistical sampling of states for proteins, which in principle should give it a key advantage. However, its top 1% TM-scores were not statistically significant higher than unperturbed AF3 for the alternative states (Figure 2A), and they were significantly lower for the preferred states (Supplementary Figure 1A). While BioEmu did not sample structures with higher top 1% TM-scores overall, it did perform better on a subset of proteins, improving/worsening the top 1% TM-scores by at least ΔTM ≥ 0.05 in 26/7 cases for the alternative states and 4/19 cases for the preferred states (Supplementary Figure 2).

To examine whether these results might be influenced by overfitting to structures in the training dataset, we filtered our dataset to proteins where both experimental structures were released after the AF3 training cut-off date (Figure 2B). While the sparsity of the dataset decreased the statistical significance of our results, the overall trend remained, with the MSA perturbing techniques improving sampling of the alternative states significantly.

If we consider the overall distribution of top 1% TM-scores for both the alternative and preferred states of each protein (Figure 2C), most of the previous conclusions remain true; unperturbed AF3 samples high TM-scores for a larger fraction of proteins compared to AF2 and all MSA perturbation techniques enhance the scores even further, with column masking achieving the highest improvements. Interestingly, BioEmu sampled a smaller fraction of protein states compared to AF3 with TM-scores above high TM-score thresholds (TM_threshold_ ≳ 0.88), but a larger fraction of proteins with TM-scores above low TM-score thresholds (TM_threshold_ ≲ 0.84).

Given the overall success of column masking, we sought to examine whether the choice of amino acid mask could influence the performance. While the standard choice [14, 16] is to mask the MSA using the unknown amino acid X, the method trivially generalizes to any other choice of amino acid. Due to the increase in computational cost to rerun the sampling for every possible amino acid mask, we limited our analysis to the 55 proteins where unperturbed AF3 sampled at least one state with a mean top 1% TM-score below 0.9. While most protein states were sampled with similar TM-scores, masking with phenylalanine (F) improved the TM-score by at least ΔTM ≥ 0.05 in 9 out of 55 cases, more than any other choice of amino acids (Figure 2D). These results do not necessarily mean that F-masking always is the best choice of amino acid column mask, nor that it performs better than X in general; in some cases, other choices of amino acids improved the TM-score significantly more than either X or F (Supplementary Figure 3), and the overall difference between X-masking and F-masking was not statistically significant when using the Wilcoxon signed-rank test. Nevertheless, these results highlight that a change of amino acid column mask improves the sampling of alternative states in some cases.

To better elucidate our findings, we examined three cases of proteins that reflected our general results in more detail.

### AF3 samples both open and closed conformations of *β*-phosphoglucomutase, whereas AF2 only samples closed conformations

*β*-phosphoglucomutase is an enzyme that upon substrate binding undergoes conformational transitions to catalyze phosphoryl group transfers. Structures of *β*-phosphoglucomutase in both open and closed conformations are present in both AF2 and AF3’s training dataset. However, all of our AF2-sampled structures had domains that were tightly in contact, closely resembling the closed state of the enzyme only (with a mean top 1% TM-scores ⟨TM_1%_⟩ = 0.99) (Figure 3), and not the open state (⟨TM_1%_⟩ = 0.83). AF3, on the other hand, sampled structures with varying degrees of inter-domain closure, matching both the closed state (⟨TM_1%_⟩ = 0.99) and the open state (⟨TM_1%_⟩ = 0.97), as well as intermediary conformations. While the top 1% of TM-scores for both states was so high for unperturbed AF3 that it could not possibly be substantially improved by the MSA-perturbing techniques, neither column masking, clustering nor stochastic subsampling worsened the scores more than ΔTM *<* 0.02 (Figure 3C).

**Fig. 3.**
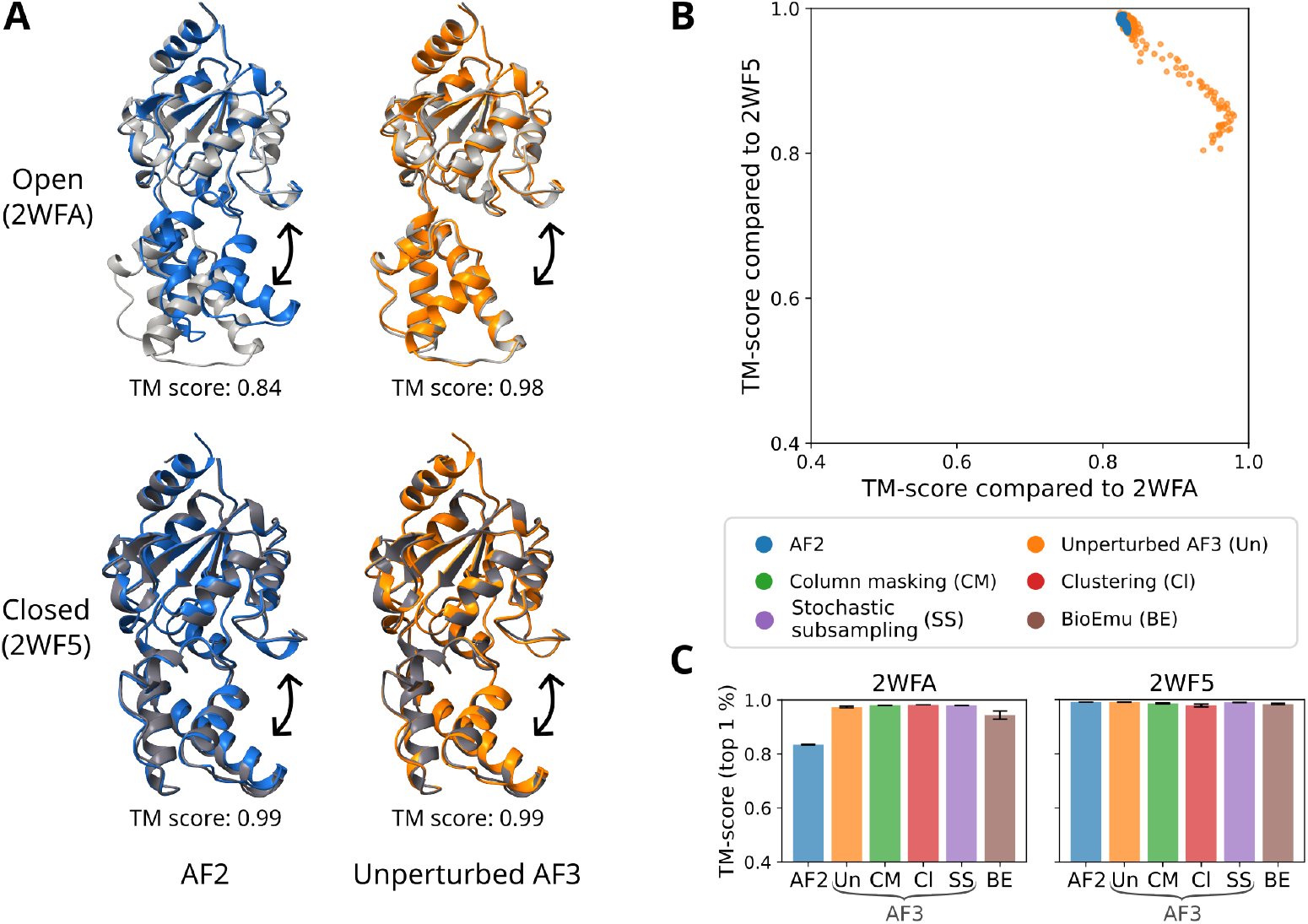
AF3 samples both open and closed conformations of *β*-phosphoglucomutase, whereas AF2 only samples closed conformations. (A) Selected structure predictions of unperturbed AF2 (blue) and unperturbed AF3 (orange) with the highest TM-score compared to an experimentally determined open state (light grey, PDB ID: 2WFA) and closed state (dark grey, PDB ID: 2WF5), respectively. The structures are visualized using ChimeraX [20]. (B) Diversity plot showing the TM-scores of all predictions to the open state and the closed state for unperturbed AF2 and unperturbed AF3. (C) Mean top 1% TM-scores for each method compared to the open and closed states. Error bars indicate the standard deviation of the top 1% TM-scores (with n=37 for clustering and n=10 for the remaining methods).

### Column masking allows AF3 to sample an additional state of a calcium-transporting ATPase

Calcium-transporting ATPases are phosporylated by ATP, which induces a conformational shift that pumps calcium ions against the concentration gradient. The entire process involves multiple conformational states, which have only recently been structurally resolved for certain subtypes. The Secretory pathway Ca^2+^-transporting ATPase type 1 is an example of this, where cryo-EM structures in multiple states were all solved after the AF3 training date cutoff [21, 22]. In our case, we found that unperturbed AF3 sampled some conformations that were highly similar to the metal-free phosphorylated (E2P) state (⟨TM_1%_⟩ = 0.95), and others that were similar to the ion-bound (CaE1) state (⟨TM_1%_⟩ = 0.95), capturing both the rearrangement and remodeling of transmembrane helices as well as the rotation of a cytosolic domain between these states. Unperturbed AF3 did not, however, sample conformations resembling the ATP and ion-bound (E1-ATP) state (⟨TM_1%_⟩ = 0.78) (Figure 4, Supplementary Figure 4). When we introduced column-masking with AF3 the sampling was broader, and also included conformations resembling the E1-ATP state (⟨TM_1%_⟩ = 0.91), with a displacement of another cytosolic domain as well as a downward movement of two transmembrane helices compared to the CaE1 state. No other tested method sampled the E1-ATP state equally well (with ⟨TM_1%_⟩ = 0.87 for clustering and ⟨TM_1%_⟩ *<* 0.83 for the remaining methods) (Figure 4C, Supplementary Figure 4).

**Fig. 4.**
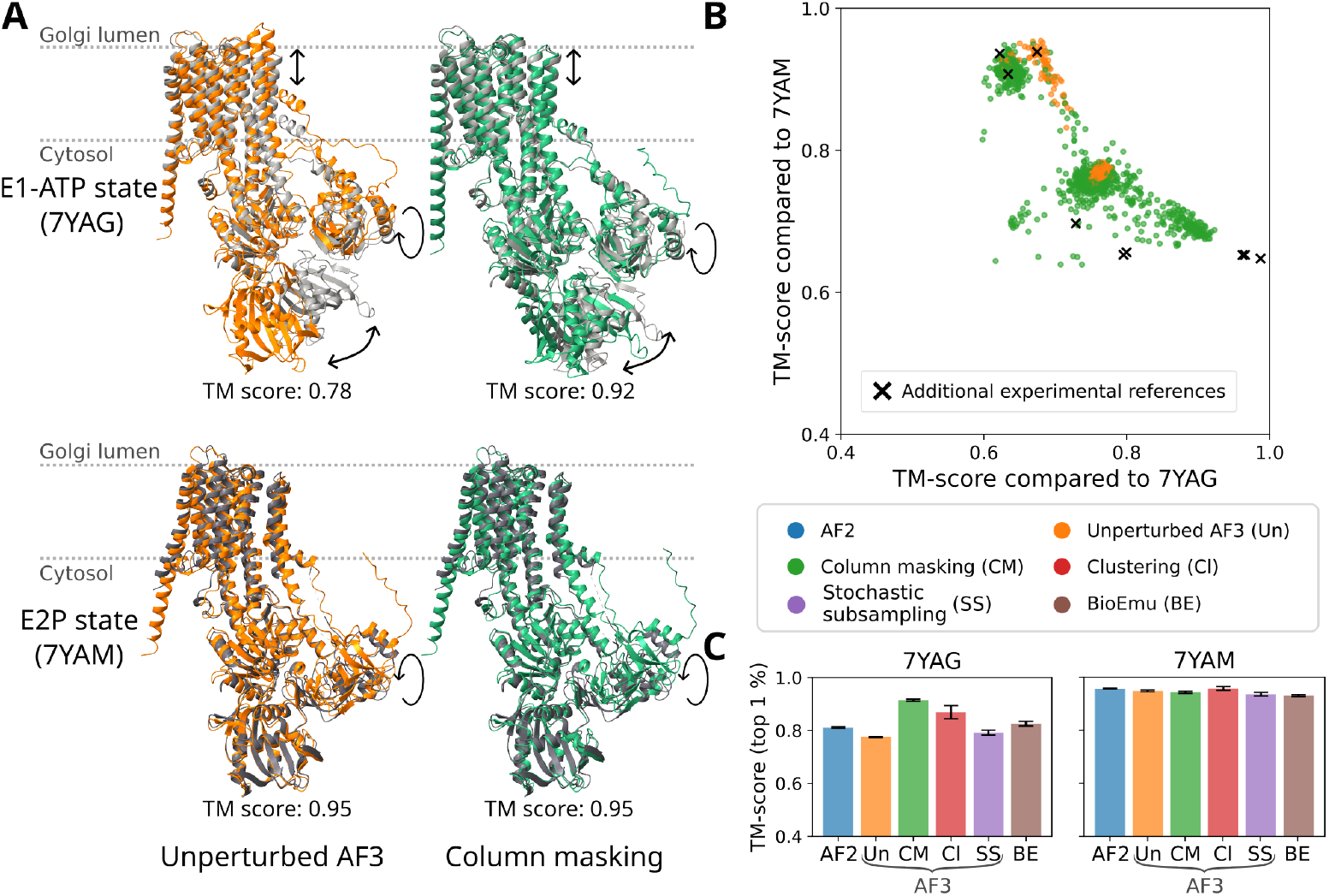
Column masking allows AF3 to sample an additional state of a calcium-transporting ATPase. (A) Selected structure predictions of unperturbed AF3 (orange) and column-masked AF3 (green) with the highest TM-score compared to reference structures in the ATP and ion-bound (E1-ATP) state (light grey, PDB ID: 7YAG) and metal-free phosphorylated (E2P) state (dark grey, PDB ID: 7YAM), respectively. The structures are visualized using ChimeraX [20]. Diversity plot showing the TM-scores of all predictions to the E1-ATP state and the E2P state for unperturbed AF3 and column-masked AF3. Crosses indicate the corresponding TM-scores for additional experimental structures when compared to the chosen reference structures. (C) Mean top 1% TM-scores for each method compared to the E1-ATP and E2P states. Error bars indicate the standard deviation of the top 1% TM-scores (with n=37 for clustering and n=10 for the remaining methods).

### Column masking by F instead of X leads AF3 to sample the apo state of an RNA helicase

In a handful of cases, the choice of amino acid mask had major influence on sampling, with especially striking difference for the Nucleolar RNA helicase 2 (Figure 5). This helicase unwinds RNA in a process that has been proposed to contain four major states: apo, pre-unwound, post-unwound/pre-hydrolysis, and post hydrolysis, where two core domains are arranged differently with respect to each other [23]. In our benchmark, the apo and post-hydrolysis states were designated as references, and none of these states were sampled well by column-masked AF3 when using X as an amino acid mask, nor by any other type of sampling method (with ⟨TM_1%_⟩ *<* 0.7 for both states and all methods) (Figure 5, Supplementary Figure 5A). Instead, it seems that all of these methods sample conformations with the core domains arranged closer to the arrangement of the post-unwound/pre-hydrolysis state, (with ⟨TM_1%_⟩ ≥ 0.97 for all methods) (Supplementary Figure 5B). However, when we masked the MSA using F instead of X, AF3 sampled additional domain arrangements, with some conformations closely resembling the apo state (with ⟨TM_1%_⟩ = 0.987). Some other choices of amino acid column masks also resulted in sampling of conformations resembling the apo state, especially D, R and W (with ⟨TM_1%_⟩ = 0.978, ⟨TM_1%_⟩ = 0.994 and ⟨TM_1%_⟩ = 0.992, respectively) (Figure 5C).

**Fig. 5.**
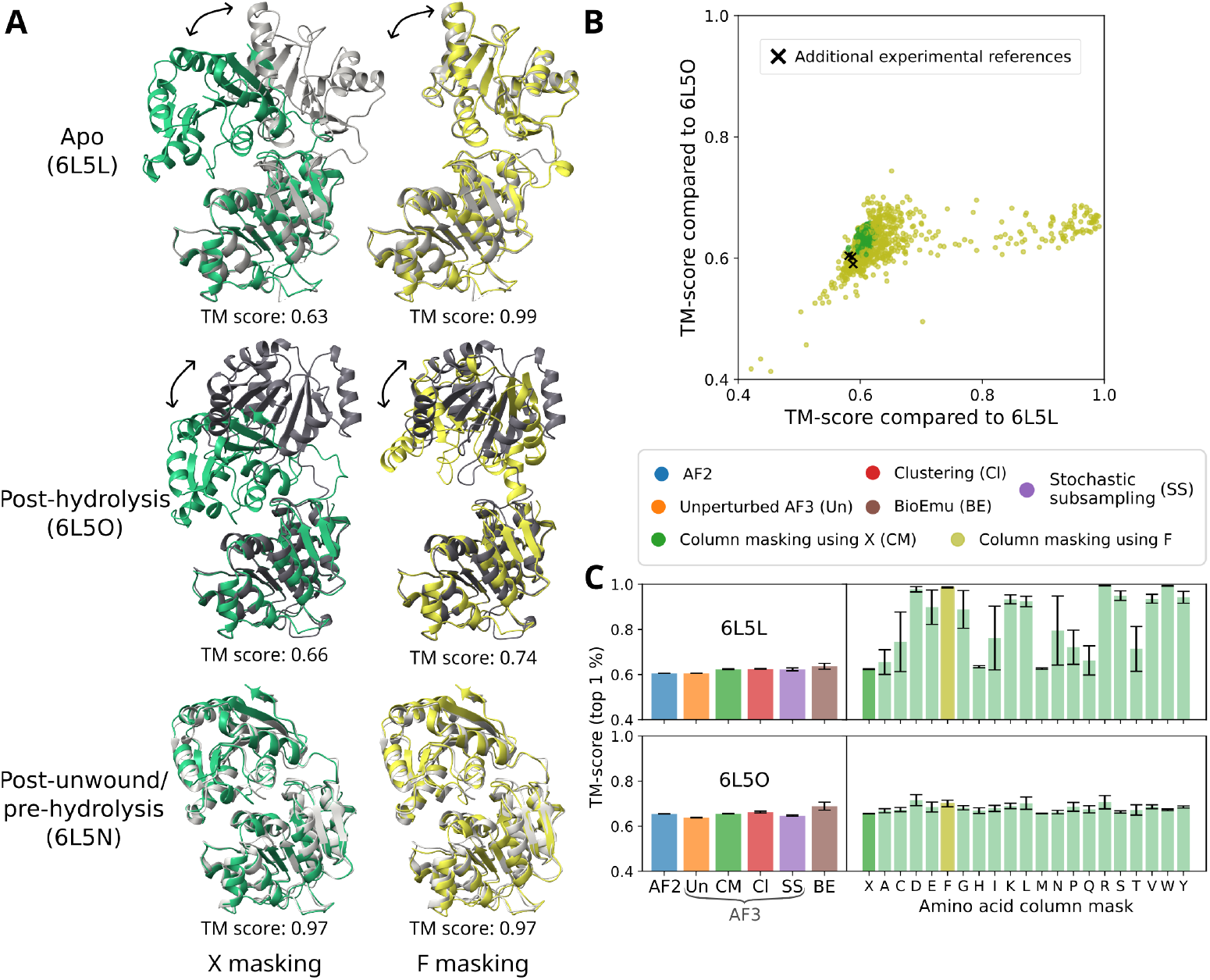
Column masking by F instead of X leads AF3 to sample the apo state of an RNA helicase. (A) Selected structure predictions of column-masked AF3 when masking using X (green) and F (yellow) with the highest TM-score compared to an experimentally determined apo state (light grey, PDB ID: 6L5L), post-hydrolysis state (dark grey, PDB ID: 6L5O), and post-unwound/pre-hydrolysis state (white, PDB ID: 6L5N). The structures are visualized using ChimeraX [20]. (B) Diversity plot showing the TM-scores of all predictions to the apo state and the post-hydrolysis state for column-masked AF3 when using either X or F as a mask. Crosses indicate the corresponding TM-scores for additional experimental structures when compared to the chosen reference structures. Mean top 1% TM-scores for each method as well as each possible choice of amino acid mask, compared to the apo and post-hydrolysis state. Error bars indicate the standard deviation of the top 1% TM-scores (with n=11 for clustering and n=10 for the remaining methods).

To ensure that these results were not statistical artifact, we resampled a total of 20,000 conformations of the Nucleolar RNA helicase 2 using X as a column mask, corresponding to the total number of conformations sampled with any other column mask than X. In this case, a single conformation had a TM-score of 0.923 compared to the experimental apo state, while the remaining conformations all had TM-scores below 0.7 (Supplementary Figure 5C).

## Discussion

Obtaining structural information for all relevant functional states of proteins is critical to understand their function. Here, we have shown that AF3 significantly improves the sampling of alternative states over AlphaFold2 on a dataset with over 100 different proteins, that AF3 performs comparably to BioEmu for the alternative states and better for the preferred states, and that MSA perturbations and particularly column masking improve the AF3 sampling further. We highlight these results in the context of *β*-phosphoglucomutase and a calcium-transporting ATPase, where AF3, in contrast to AF2, samples an open state of the enzyme, and where column-masked AF3 unlike unperturbed AF3 samples the ATP and ion-bound state of the transporter. These results extend those of the recent AFsample3 study [16], which showed that column masking in AF3 improved such sampling on the Cfold dataset [24], that overlaps with our dataset for only 17 of the 107 tested proteins. We have further shown that a change of amino acid column mask, although not generally better, improves the sampling of alternative states in some cases.

We cannot exclude the possibility that the improved sampling upon change of amino acid column mask at least in part may be a result of overfitting to training data. However, since the switch of amino acid mask also improved the top 1% TM score even when the experimental reference was released after the AF3 training date cutoff (Supplementary Figure 3), it is unlikely to fully account for the improvement. Possibly, the amino acid-masked columns can act as conserved residues that AF3 has learned to place in specific environments, since conserved residues may cluster at specific sites on the proteins [25–27]. Nonetheless, using different column masks can improve the sampling of some protein states over simply generating more samples using the same mask, as clearly seen for the apo state of the Nucleolar RNA helicase 2. Practically, our results suggest that column-masked AF3 is a favorable method for predicting proteins in multiple conformations, and that it could be beneficial to change the amino acid mask, e.g. to F, if the standard X mask fails.

While the MSA-perturbing methods and BioEmu did sample conformations that were highly similar (with TM-scores above 0.9) to both experimental reference structures for more than half the proteins in our data set, for more than a quarter of our tested proteins all methods failed to sample structures with TM-scores above 0.8 to at least one experimental reference structure (Figure 2A). Thus, while the methods generally improve predictions, they are limited in their predictive capabilities and struggle in a significant number of cases even when we know there are at least two different states sampled. Reliably identifying multiple biologically relevant conformations *a priori* even for cases where there is no such biological information would be significantly more challenging, and still appears to be beyond the general capability of AI-based methods.

In assessing the different methods, we have compared only the top 1% TM-scores to the given reference structures. It is possible that some methods may improve sampling of the alternative states but also introduce noise and predict many non-physiological states. This introduces the challenge of distinguishing physiological relevant structures from sampling artifacts. One option to resolve this is to cluster the structures and use confidence scores to identify the relevant conformations, which has been shown to work well for column masking [16]. A complementary approach is to use the predicted models as a prior and identify the relevant conformations using functional and/or low resolution experimental data, either starting from the total ensemble of AlphaFold-sampled structures [6, 7], or by modifying the diffusion step in AF3 to directly guide the sampling, which has been shown to work for high-resolution X-ray and Cryo-EM data [28, 29]. Given that the MSA perturbations enhance the sampling of alternative states per fixed latent space, they might also be useful for simultaneously guiding multiple samples in the diffusion model to low-resolution experimental data which contain information on multiple states.

In conclusion, we have shown how several different MSA perturbations improve sampling of alternative states in AlphaFold3, and there are significant differences in their performance. Although these methods alone do not capture the entire underlying Boltzmann distribution, they provide highly valuable structural hypotheses, and a method specifically trained to generate Boltzmann distributions of conformations did not perform better. By complementing other computational methods and experimental data, these predictions enable a richer understanding of the underlying biological processes, and the conclusions will hopefully be of use to improve training of methods more specifically designed to sample full conformational landscapes.

## Methods

### Dataset

Monomeric experimental protein structures were selected from the OC23 [14] and IOMemP [30] datasets, as well as the BioEmu multi-conformation benchmark dataset v0.1 [17] for the proteins that had two experimentally solved distinct states. Three out of the 110 unique proteins were excluded due to structure prediction failures at inference time. The combined data set (Supplementary Table 1, Supplementary Figure 6) includes soluble proteins in open and closed states, membrane proteins in inward and outward-facing states, proteins with cryptic pockets in apo and holo states as well as proteins with domains that move between states.

### Structure predictions

For all proteins, structure predictions were performed using the canonical sequences from UniProt [31]. MSAs were generated for each protein using the default data pipeline implemented in AlphaFold3 [15], which utilizes JackHMMER [32] and HHBlits [33] searches on UniRef90 [34], UniProt [31], Uniclust30 [35], BFD [36] and MGnify [37].

Exactly 1000 predictions were generated for each protein and method except for MSA clustering, where at least 1000 structure predictions were generated (details below). AF2 predictions were generated using AlphaFold v2.3.2 [8] in ColabFold v1.5.5 [38] and BioEmu predictions were generated using bioemu-v1.1 [17] with `--filter_samples=False`, both methods using the unpaired MSAs from the AF3 data pipeline as input. All AF3 structure predictions were generated using AlphaFold v3.0.0 with 5 samples per random seed. For unperturbed AF3, inference was run for 200 random seeds. For stochastic subsampling, the `num_msa` parameter was restricted in the evoformer configuration from the default value of 1024 to a value in `num_msa` ∈ *{*1, 2, 4, 8, 16, 32, 64, 128, 256, 512*}*, similarly to what was done for AF2 [12], and inference was run for 20 random seeds per value of `num_msa`. For column masking, columns were stochastically masked in the MSA following the approach used for AF2 [14] and recently for AF3 [16], and the fraction of masked columns *f* was varied from 0.1 to 1.0 in increments of 0.1, with 20 random seeds per masking fraction. For MSA clustering, the MSA clustering script provided with the AF-Cluster study was used[13] to cluster the unpaired MSA, and inference was run on all clustered MSAs separately, discarding the paired MSA. To ensure at least 1000 predictions per protein across *n*_clust_ clustered MSAs, inference was performed using *n*_seeds_ = ⌈200*/n*_clust_⌉ random seeds for each clustered MSA.

### Statistical analysis

Statistical significance of the difference in performance between any pair of methods was assessed using the Wilcoxon signed-rank test implemented in the SciPy library [39] in Python.

## Declarations

## Acknowledgements

AlphaFold runs were performed using the computing facilities of the Berzelius through NAISS (grant no. Berzelius-2025-269). This work was funded by grants from the Swedish Research Council (VR; 2019-02433, 2021-05806), the Knut and Alice Wallenberg foundation (KAW; 2023.0254) and the BioExcel-3 Centre-of-Excellence (EuroHPC Joint Undertaking; 101093290) to E.L.

## Data and code availability

The code used for generating all structure predictions, analyzing the data, and recreating the figures, along with the raw TM-scores of all predictions and the top 1% scoring conformations of the highlighted proteins, are available at https://doi.org/10.5281/zenodo.19336934.

## Author contributions

**Conceptualization:** S.E.L. **Data curation:** S.E.L. **Formal analysis:** S.E.L., I.N., J.K.A. **Investigation:** S.E.L., I.N., J.K.A. **Methodology:** S.E.L. **Project administration:** R.J.H., E.L. **Resources:** S.E.L. **Software:** S.E.L. **Supervision:** E.L., R.J.H., **Validation:** S.E.L. **Visualization:** S.E.L. **Writing – original draft:** S.E.L. **Writing – review & editing:** S.E.L, E.L., R.J.H.

## Competing interests

The authors declare no competing interests.

## Notes

https://zenodo.org/records/19336934

### Competing Interest Statement

The authors have declared no competing interest.

