## Supplementary Information for "Several multiple sequence alignment perturbation methods enhance AlphaFold3 sampling of alternative protein states"

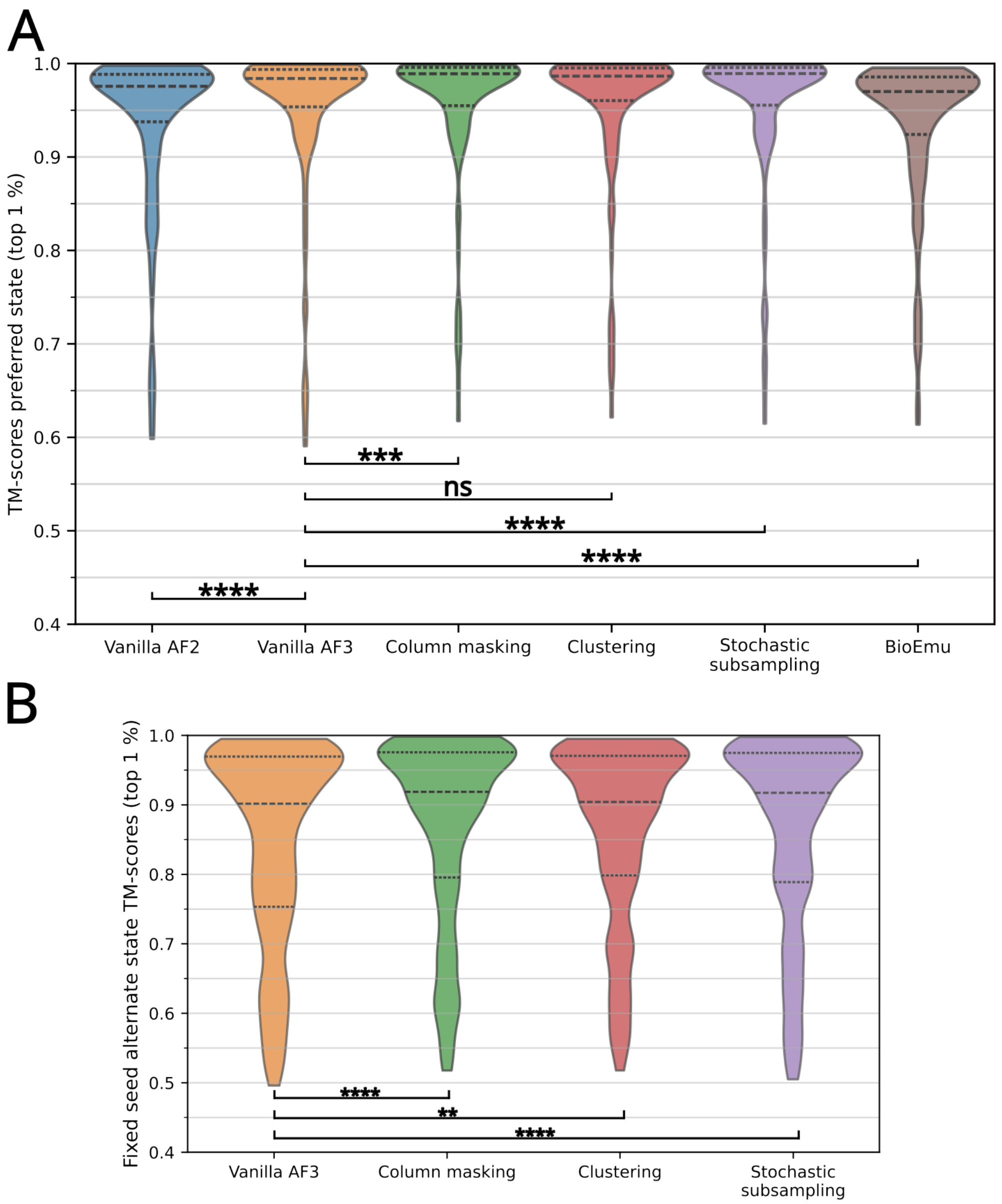


**Supplementary Figure 1 Additional TM-score distributions.** Violin plots showing the distributions of the mean top 1% TM-scores of (A) the preferred state for all methods, and (B) the alternative state per latent space in AF3 (i.e. per fixed seed), both across the entire dataset of 107 proteins. Statistical significance is assessed using the Wilcoxon signed-rank test (*: $p < 0.05$, **: $p < 0.01$, ***: $p < 0.001$, ****: $p < 0.0001$).

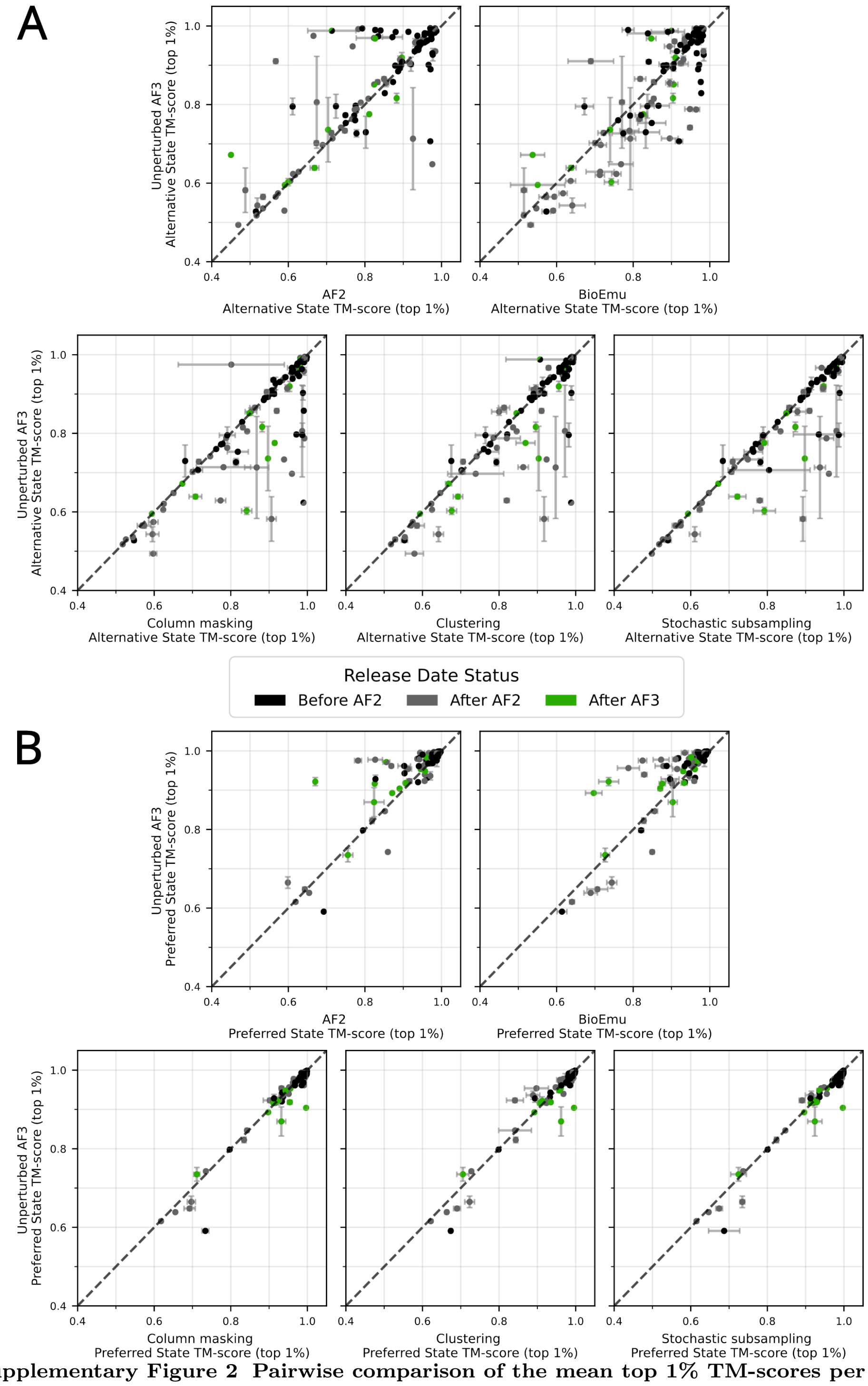


**Supplementary Figure 2 Pairwise comparison of the mean top 1% TM-scores per protein between unperturbed AF3 and the remaining methods.** Shown for (A) the alternative state and (B) the preferred state. Error bars indicate the standard deviation of the corresponding top 1% TM-scores.

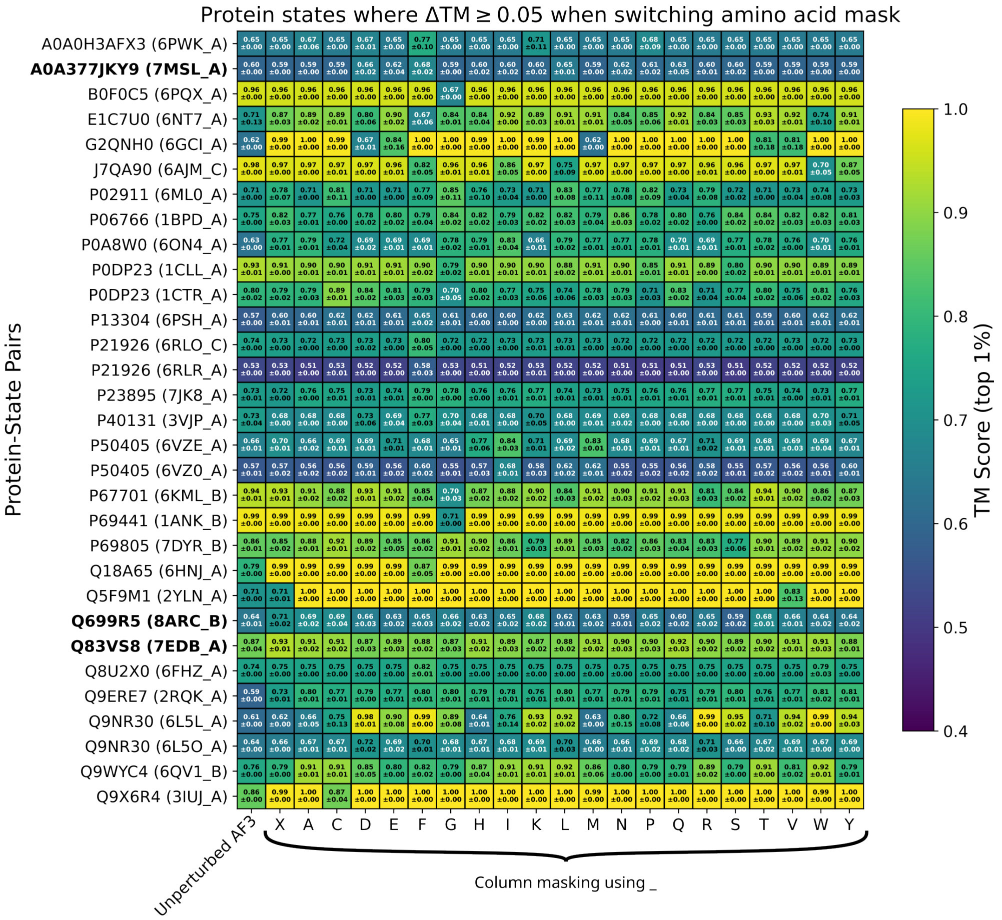


**Supplementary Figure 3 The optimal choice of amino acid column mask varies for different proteins.** Heatmap of the top 1% TM-scores (mean ± standard deviation) for either unperturbed AF3 or column-masked AF3 with a given mask, for the 31 out of 110 protein-state pairs (representing both states across the 55 evaluated proteins) where the mean top 1% TM-score differed by $\Delta\text{TM} \geq 0.05$ when switching from X to any other amino acid mask. Reference structures that were released after the AF3 training date cutoff are shown in bold.

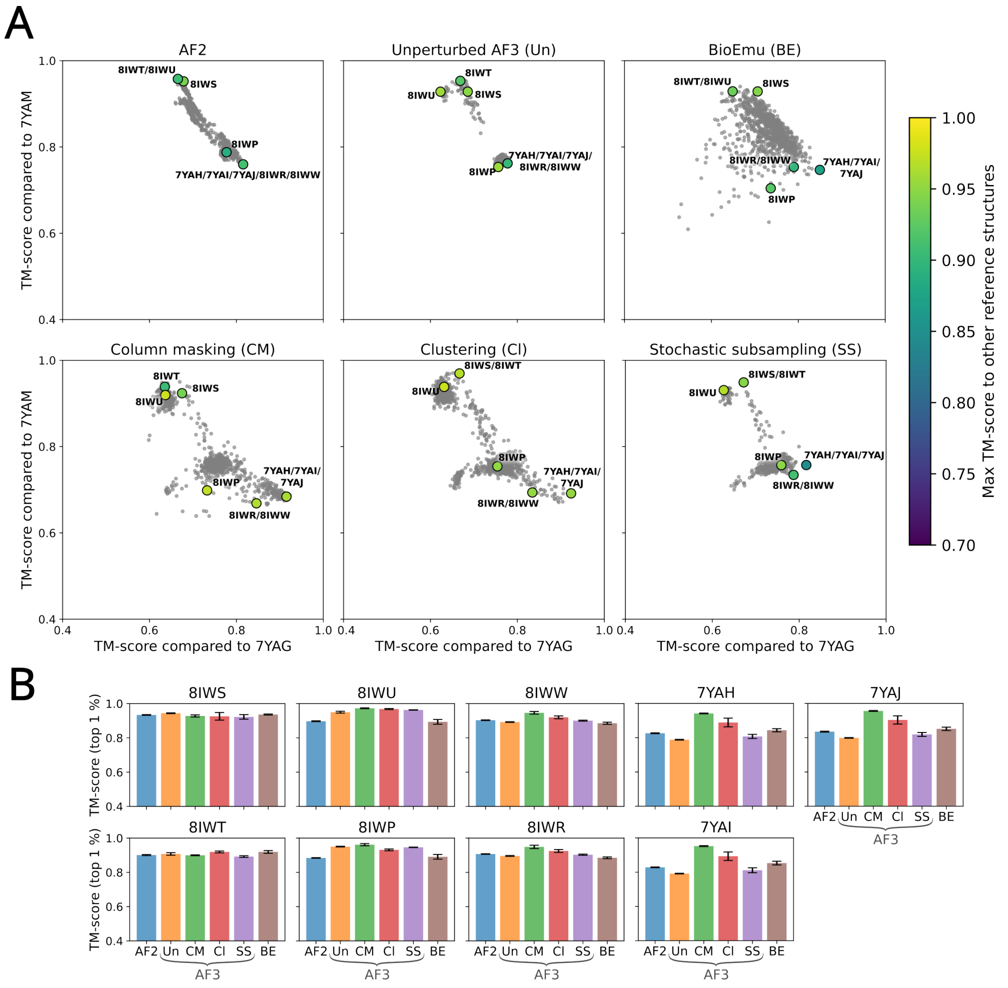


**Supplementary Figure 4 Structure predictions of the Secretory pathway $Ca^{2+}$-transporting ATPase type 1 for all methods and all PDB-deposited reference states.** (A) Diversity plots showing the TM-scores of all predictions to reference structures in the E1-ATP state (PDB ID: 7YAG) and the E2P state (PDB ID: 7YAM). Colored points mark the structure predictions that are closest to another reference structure in the PDB. The 8IWP reference structure represents an ion-bound (CaE1) state. (B) Mean top 1% TM-scores for each method compared to all additional PDB-deposited structures. Error bars indicate the standard deviation of the top 1% TM-scores (with $n = 37$ for clustering and $n = 10$ for the remaining methods).

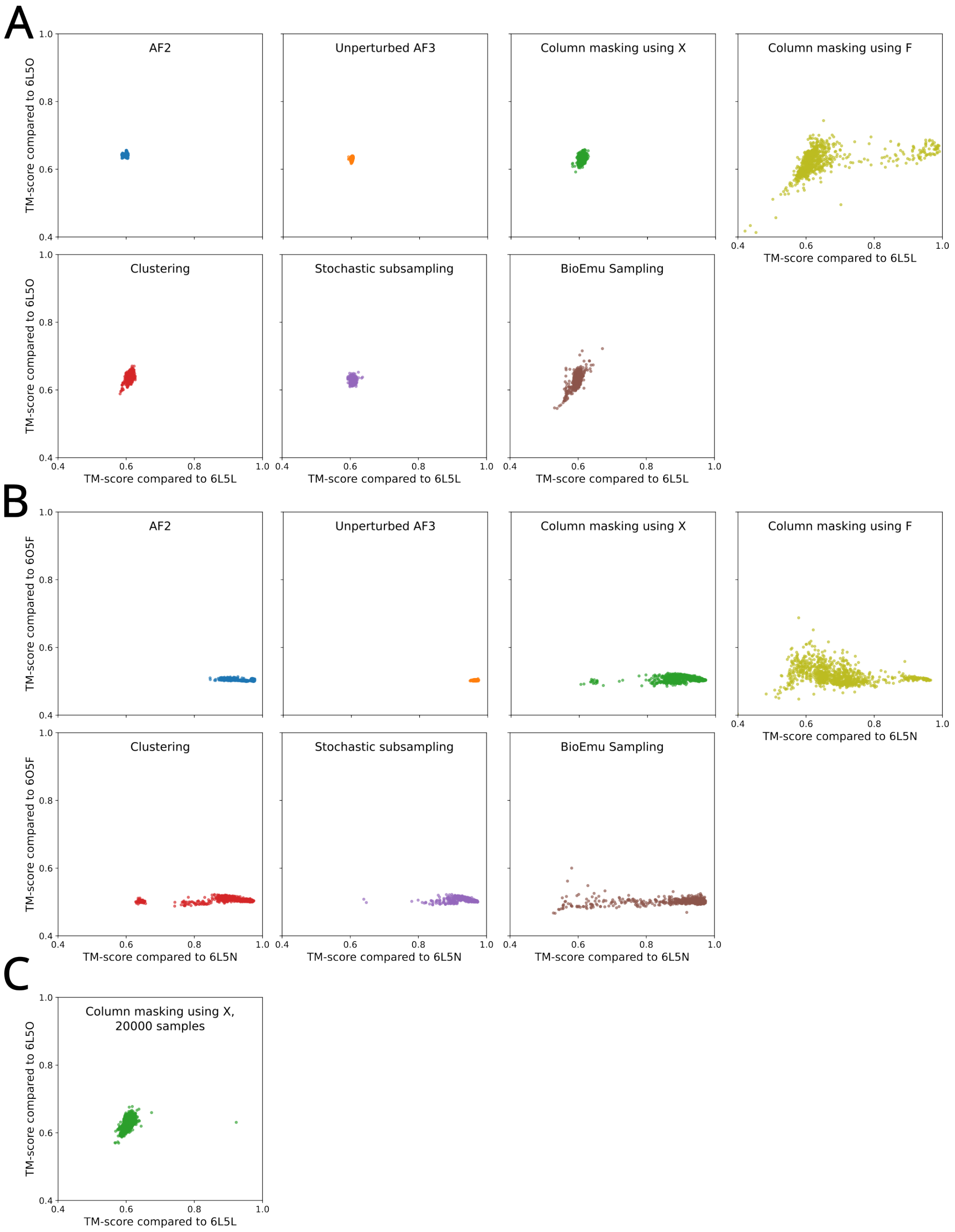


**Supplementary Figure 5 Structure predictions of the Nucleolar RNA helicase 2 for all methods and all PDB-deposited reference states.** Diversity plots showing the TM-scores for all predictions to experimental reference structures. Predictions are shown from (A, B) all methods, and (C) an expanded rerun of column masking. The reference structures corresponds to (A, C) the apo state (PDB ID: 6L5L) and the post-hydrolysis state (PDB ID: 6L5O), and (B) the post-unwound/pre-hydrolysis state (PDB ID: 6L5N) and the pre-unwound state (PDB ID: 6O5F).

**Supplementary Table 1** Proteins and reference structures used in the benchmark.

| UniProt ID | PDB ID 1 | PDB ID 2 |
|---|---|---|
| A0A075Q0W3 | 6MKA_A | 6MKJ_A |
| A0A0H3AFX3 | 6PWJ_A | 6PWK_A |
| A0A377JKY9 | 7F7P_A | 7MSL_A |
| A0QTT2 | 7CY2_A | 7CYR_A |
| A2RJ53 | 3FTO_A | 3DRF_A |
| A6UVT1 | 6HAC_A | 6HAE_A |
| B0F0C5 | 6PQN_A | 6PQX_A |
| B3EYN2 | 5HO2_A | 5HO0_A |
| B7IE18 | 6NC7_A | 6NC6_A |
| E1C7U0 | 6NT7_A | 6NT6_A |
| F1NCD6 | 7MJS_X | 7N98_A |
| G2QNH0 | 6GCI_A | 4C9J_B |
| J7QA90 | 6GTS_B | 6AJM_C |
| J7QAK3 | 6E9N_B | 6E9O_B |
| K0JNC6 | 7OZ3_A | 7B5Y_A |
| M1GSK9 | 6P3X_A | 7UU3_A |
| M1VAN7 | 6A6N_A | 7DQV_A |
| O74933 | 2YQC_A | 2YQS_A |
| O76728 | 4BP8_A | 4BP9_A |
| O86150 | 5C78_A | 6HRC_D |
| P00558 | 2XE6_A | 2WZD_A |
| P00766 | 2CGA_B | 1AFQ_C |
| P02787 | 1BP5_A | 1RYO_A |
| P02911 | 6ML0_A | 6MLP_A |
| P02925 | 1URP_A | 2DRI_A |
| P03047 | 7UBJ_A | 7UBL_A |
| P03960 | 7LC3_B | 7BGY_B |
| P06766 | 1BPD_A | 2BPG_E |
| P08037 | 1PZT_A | 1PZY_D |
| P08709 | 1JBU_A | 1WUN_B |
| P0A7D4 | 1ADE_A | 1CIB_A |
| P0A7F3 | 2AIR_D | 1ZA1_D |
| P0A8V6 | 1E2X_A | 1H9G_A |
| P0A8W0 | 6ON4_A | 6WFQ_A |
| P0AEQ3 | 1WDN_A | 1GGG_A |
| P0AEX9 | 3PUW_A | 1FQC_A |
| P0AEY8 | 6VS1_A | 6GV1_A |
| P0AG16 | 1ECJ_D | 1ECC_B |
| P0CL43 | 6XFJ_A | 6XFL_A |
| P0DP23 | 1CLL_A | 1CTR_A |
| P12758 | 1K3F_B | 1U1D_F |
| P13304 | 6PX4_A | 6PSH_A |
| P18238 | 6GCI_A | 4C9J_B |
| P19491 | 1MY1_C | 1FTL_A |
| P20789 | 6Z4Q_A | 6Z8N_A |
| P21589 | 7QGA_A | 4H2I_A |
| P21926 | 6RLO_C | 6RLR_A |
| P23847 | 1DPE_A | 1DPP_A |
| P23895 | 7MGX_A | 7JK8_A |
| P24182 | 1BNC_B | 2V5A_A |
| P26281 | 1HKA_A | 3IP0_A |
| P28366 | 1M74_A | 1TF2_A |
| P30878 | 7L17_A | 4M64_A |
| P31133 | 6YED_A | 6YE0_A |
| P33284 | 3O80_A | 3O8M_A |
| P37487 | 1K23_C | 1WPM_B |
| P40131 | 3TEE_A | 3VJP_A |
| P43005 | 8CTC_A | 6S3Q_A |
| P44542 | 2CEY_A | 6H76_A |
| P45568 | 1K5H_C | 2EGH_B |
| P50405 | 6VZE_A | 6VZ0_A |
| P54708 | 8IJL_A | 7X23_A |
| P56817 | 1W50_A | 3IXJ_C |
| P60338 | 1EXM_A | 1HA3_B |
| P60752 | 7SEL_A | 8DMM_A |
| P61586 | 1XCG_B | 1OW3_B |
| P62495 | 2KTV_A | 2KTU_A |
| P62593 | 1JWP_A | 1PZO_A |
| P67701 | 6JQ4_A | 6KML_B |
| P68082 | 7DGJ_A | 6LS8_A |
| P69441 | 4AKE_B | 1ANK_B |
| P69801 | 7DYR_A | 6K1H_B |
| P69805 | 7DYR_B | 6K1H_A |
| P71447 | 2WFA_A | 2WF5_A |
| P79345 | 1NEP_A | 2HKA_C |
| P84080 | 1RRG_A | 1S9D_A |
| P98194 | 7YAG_A | 7YAM_A |
| P9WPY3 | 2IYT_A | 2IYQ_A |
| Q02153 | 6JT0_B | 7D9R_B |
| Q02763 | 1FVR_A | 2OO8_A |
| Q16539 | 2ZB1_A | 3HL7_A |
| Q16658 | 3P53_A | 6I11_A |
| Q18A65 | 6HNJ_A | 6HNI_A |
| Q27686 | 1PKL_B | 3HQP_P |
| Q29495 | 1B6B_A | 1KUV_A |
| Q2FYI5 | 8E2C_A | 8E2B_A |
| Q31PX7 | 7N82_A | 6UF2_A |
| Q53W80 | 7C63_A | 7C66_A |
| Q58L87 | 2WGB_A | 2V57_A |
| Q5F9M1 | 3ZSF_A | 2YLN_A |
| Q63GK8 | 6BVG_A | 5IWS_A |
| Q699R5 | 8ARC_B | 8ARB_B |
| Q6MLJ0 | 6BTX_A | 5AYM_A |
| Q72J05 | 5MKK_B | 6RAJ_B |
| Q7DAU8 | 3L6G_A | 3L6H_A |
| Q83VS8 | 7E8R_A | 7EDB_A |
| Q8U2X0 | 6FHZ_A | 3VVS_A |
| Q8WSF8 | 3PEO_G | 2BYS_J |
| Q9DA75 | 7MJS_X | 7N98_A |
| Q9ERE7 | 2RQM_A | 2RQK_A |
| Q9NR30 | 6L5L_A | 6L5O_A |
| Q9SS90 | 6K8B_A | 6K85_B |
| Q9WYC3 | 3QF4_A | 6QV1_C |
| Q9WYC4 | 4Q4A_B | 6QV1_B |
| Q9X6R4 | 3IUJ_A | 3IUQ_A |
| Q9X9P9 | 2OLO_A | 2OLN_A |
| Q9Z4N6 | 1SI1_A | 1SI0_A |

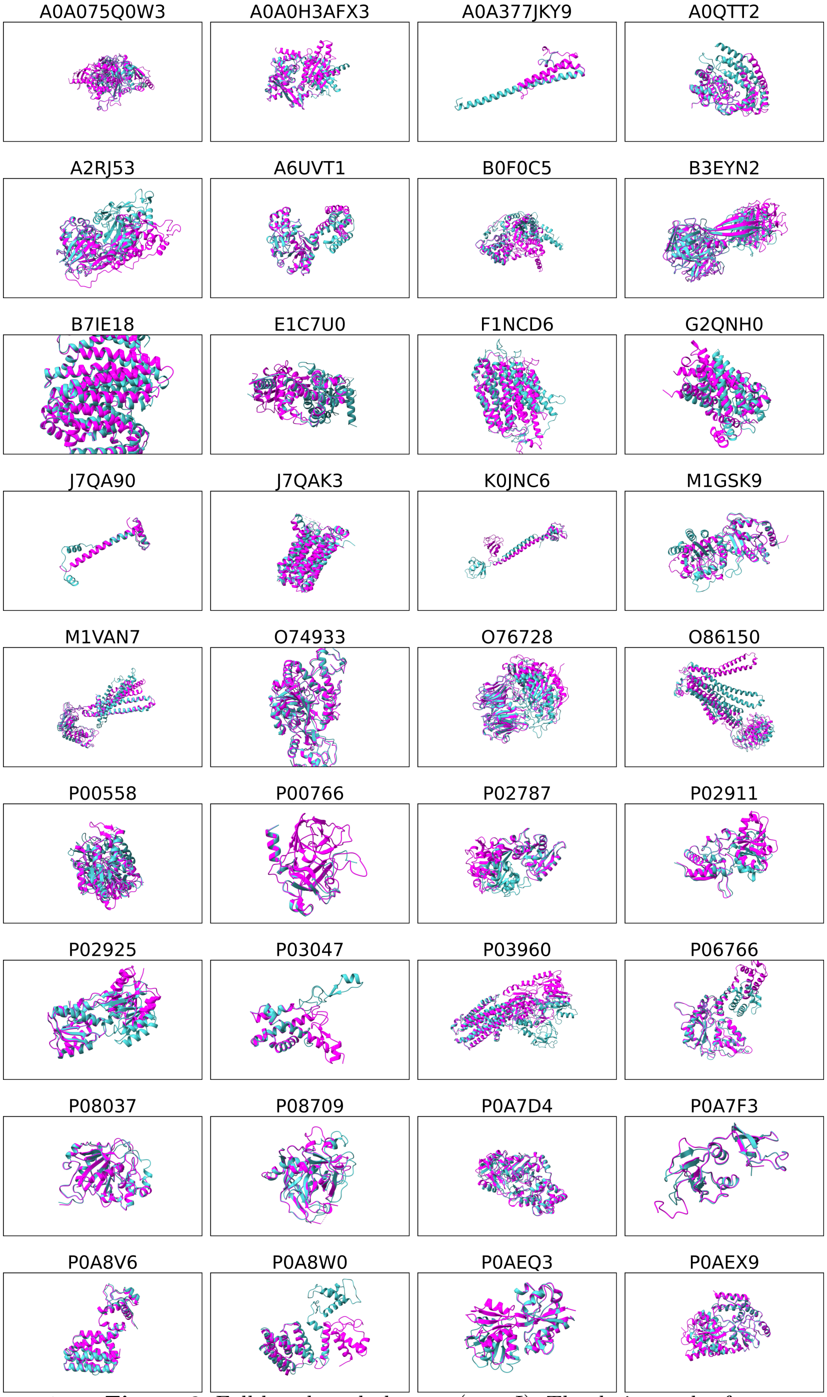


**Supplementary Figure 6** Full benchmark dataset (part I). The designated reference states are visualized for each UniProt ID using ChimeraX [20].

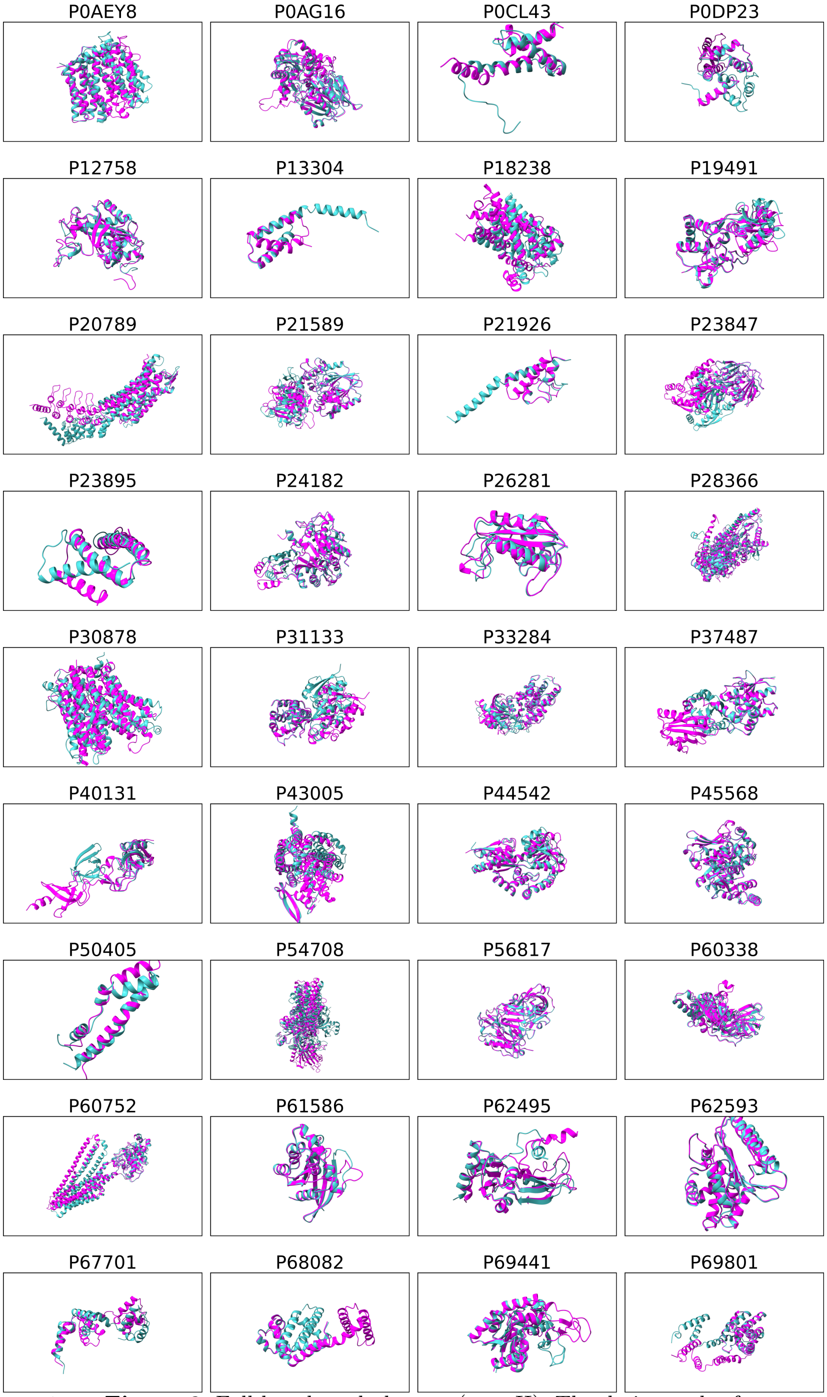


**Supplementary Figure 6** Full benchmark dataset (part II). The designated reference states are visualized for each UniProt ID using ChimeraX [20].

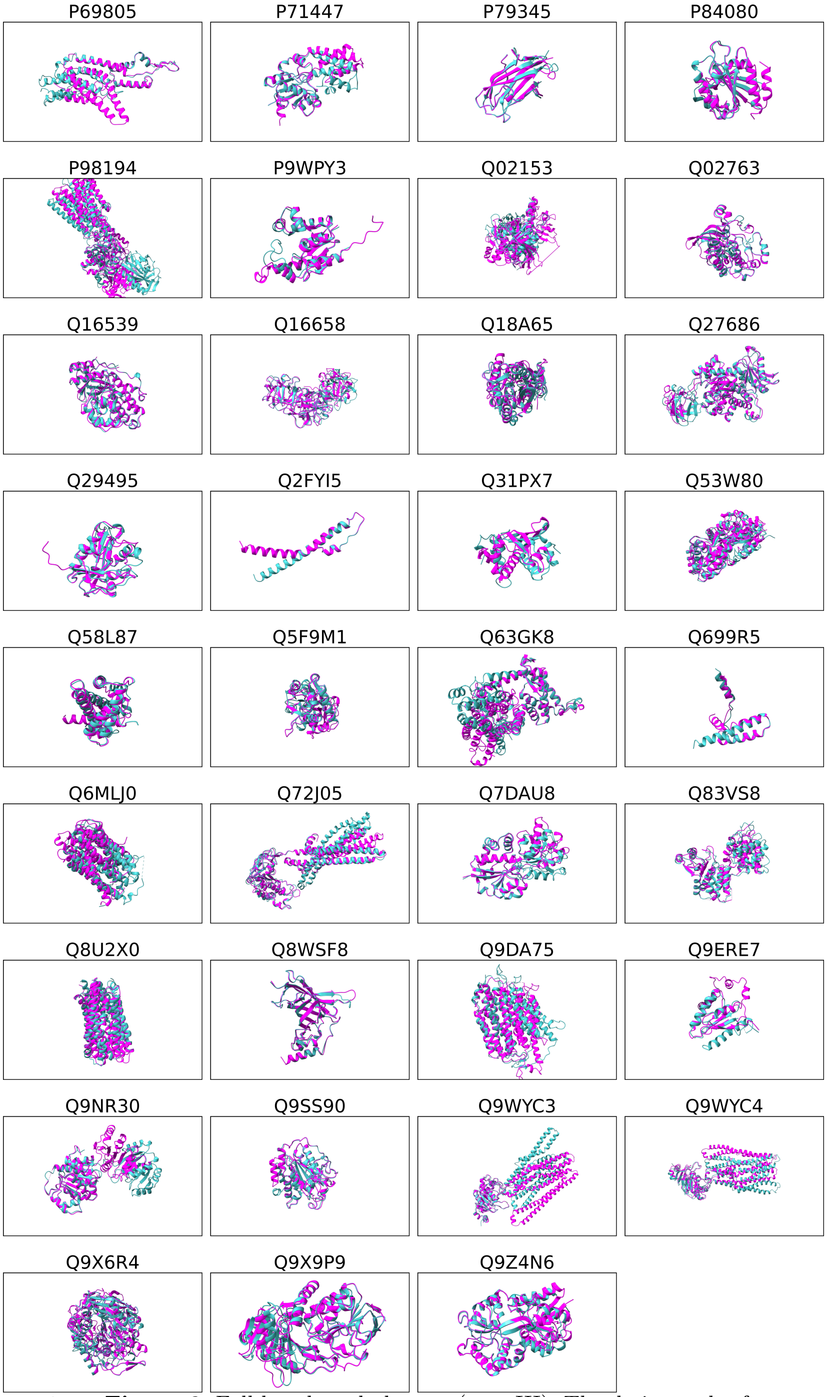


**Supplementary Figure 6** Full benchmark dataset (part III). The designated reference states are visualized for each UniProt ID using ChimeraX [20].
